# Identifying drivers of spatio-temporal variation in survival in four blue tit populations

**DOI:** 10.1101/2021.01.28.428563

**Authors:** Olivier Bastianelli, Alexandre Robert, Claire Doutrelant, Christophe de Franceschi, Pablo Giovannini, Anne Charmantier

## Abstract

In a context of rapid climate change, the influence of large-scale and local climate on population demography is increasingly scrutinized, yet studies are usually focused on one population. Demographic parameters, including survival, can vary significantly across populations of a given species, depending on global or local climatic fluctuations but also on many other population-specific parameters such as breeding density, habitat naturalness, predation or parasitism. Such ecological differences between populations could lead to different paces-of-life (POL), whereby populations where individuals display higher reproductive investment and bolder behaviours would have lower survival probabilities. We use here long-term (19 to 38 years) monitoring datasets from four Mediterranean populations of blue tits (*Cyanistes caeruleus*) to investigate the effects of sex, age class, large-scale and local climate temporal variation and population breeding density, on adult survival, using Capture-Mark-Recapture modelling. Environment heterogeneity in these four populations (two in evergreen and two in deciduous forests) has been linked to strong multi-trait phenotypic variation, suggesting blue tits in deciduous forests display faster POL compared to their conspecifics in evergreen habitats. The present results show heterogeneity in average survival probabilities across the four populations, with, as predicted, lower survival in the ‘fast’ blue tits occupying deciduous habitats. Interestingly, the year-to-year variation in survival probabilities was synchronous among populations. This suggests that regional environmental conditions could drive survival fluctuations across populations. However, breeding densities were not correlated across populations, and we found no evidence that adult survival is correlated with either large-scale or local, climate temporal variation in these four blue tit populations. Finally, two of the focal populations displayed a linear temporal decrease in adult survival over the study period, calling for further investigation to explain this decline. Overall, this multi-site study shows that blue tit parental survival from one spring to the next can vary substantially across years, in a synchronous way across populations, yet the climate indices we used are not correlated with the temporal variation. This calls for further investigations in other potential drivers such as resource (in particular insect) abundance, predation or parasitism.

## Introduction

Many temporal variations in ecological systems can be decomposed in cycles (e.g. daily, seasonal, multi-annual), in longer term trends, and in remaining ‘noise’ (e.g. year-to-year variation) (Wolkovich et al., 2014, André & Rousset, 2020). These variations often result from abiotic environmental changes over time, such as climate or local weather variations, which in turn result in biotic responses to these changes, e.g. morphological, behavioural, physiological, phenological and/or demographic variations at the population scale. Multiple studies have shown that climate can influence numerous biological processes and biodiversity patterns, with ecological and evolutionary consequences (e.g. Norberg et al., 2012, Woodbridge et al., 2021). In particular, the recent ecological literature informs us that meteorological year-to-year variation can influence population demography in plants (Chang-Yang et al., 2016, Dalgleish et al., 2011) and animals (Selonen et al., 2016, Wood et al., 2016), while recent trends in climate change cause temporal trends in demographic components and their variance across many different taxa of plants (Williams et al., 2015) and animals (Massardier-Galata et al., 2017), including birds (Alves et al., 2019, Gamelon et al., 2017). For example, in polar bears, survival of cubs to recruitment is highly dependent on their mother’s body condition in autumn, which itself depends on weather conditions, while the population demography of this species is highly impacted by the increasing reduction of sea ice availability, resulting in strong conservation concerns (Laidre et al., 2020). However, the demographic consequences of climate variation, and the link between climate, ecological factors, and demographic effects, are still insufficiently explored (see reviews Chevin et al., 2013, Visser & Gienapp, 2019).

Recent studies have shown that population density can play a major role in the impact of climate (and of climate-induced changes in traits) on population dynamics (Gamelon et al., 2017). In the blue petrel *Halobaena caerulea* for example, population crashes occur in years with both poor conditions and high densities (Barbraud & Weimerskirch, 2003). McLean and colleagues have argued that species or populations with strong density-dependent effects on population dynamics will have more robust demographic rates (*i.e.* survival and fecundity) when facing strong climate fluctuations because density-dependent processes can buffer negative demographic consequences of climate change (McLean et al., 2016). An elegant example of such buffering effect of density comes from a study of Dutch great tits *Parus major*, where warmer springs result in a detrimental mismatch between the bird breeding phenology and their main prey seasonal peak (Visser et al., 2006). While this phenology mismatch has fitness consequences for the birds, such that spring warming translates into stronger selection for earlier breeding, an increased mismatch does not result in decreased population growth (Reed et al., 2013) because of density-dependent regulation. During warm springs, great tits have a reduced breeding success, yet their fledglings show increased survival due to relaxed competition. Such examples highlight the importance of considering density-dependent effects when exploring demographic consequences of climate change.

In a context of large-scale rapid climate change, it is also important to determine whether the meteorological and climatic variations with which populations are (and will be) confronted are likely to have a similar impact on their demography, depending on their location. The spatial synchrony of demographic parameters and local population dynamics (Robert, 2009), or on the contrary their divergence (Cuervo & Moller, 2013), is a key element of species dynamics on a large spatial and temporal scale (Siriwardena et al., 1998). Demographic parameters, including survival, can vary significantly across populations of a given species, depending for instance on large scale climatic fluctuations (Post & Stenseth, 1998, Mazerolle et al., 2005), local climatic conditions and resource availability (Winkler et al., 2014, Senner et al., 2017), predation and parasitism (Watson, 2013, DeCesare et al., 2014), interspecific competition (Gustafsson, 1987), and many factors related to human activities (Hõrak & Lebreton, 1998, Porneluzi & Faaborg, 1999, Cartwright et al., 2014). Variation in survival across populations can also arise because of differences in population age-structure or sex ratio (Loison et al., 1999, Clutton-Brock & Isvaran, 2007). A previous study by Grosbois and colleagues analyzed blue tit survival in three Mediterranean blue tit (*Cyanistes caeruleus*) populations. Results from this study indicated that adult survival differed considerably both among years and among populations, and that the pattern of interannual variation in survival was similar among populations, suggesting that adult survival in these blue tit populations was influenced by environmental factors, such as climate, that operate at a relatively large spatial scale. In particular, adult survival was correlated with both local-scale weather conditions (summertime and wintertime index combining rainfall, temperature and wind variables) and a large-scale tropical index in early summer: rainfall in the Sahel. The authors noted that the Sahel rainfall index could represent either a tropical influence on European weather, or be related to local climate in a way that is not captured by their local summer climate index. Overall, while there are reasons to expect parallel variation in survival between populations of the same species as found in the blue tit study, it is also likely that it will not be the case if large scale climatic factors have a minor influence compared to local factors.

Here, using a long-term monitoring dataset on four Mediterranean populations studied across 19 to 38 years, we investigated the effects of sex, age class (one year old breeders *versus* older), large-scale and local climate temporal variation and population breeding density, on the survival of breeding adults in a temperate passerine bird, the Blue tit. This small passerine is a non-migratory, seasonal, hole-nesting breeder, weighing around 11g on the mainland and less than 10g in Corsica (smaller sub-species *C. c. ogliastrae)* (Charmantier et al., 2016), with female-biased dispersal (Garcia-Navas et al., 2014). Blue tits are short-lived, with recruitment rates of typically 5-20%, a mean inter-annual survival rate close to 50% and a mean life expectancy of 2 years on average (Garcia-Navas et al., 2014, Hadfield et al., 2006, Lambrechts et al., 2004). Survival in adult blue tits has been previously related to many dimensions of individual variation, such as pair fidelity (Culina et al., 2015), individual heterozygosity (Olano-Marin et al., 2011), immigrant *versus* resident status (Garcia-Navas et al., 2014), reproductive effort and parasitism (Stjernman et al., 2004), body mass (Nord & Nilsson, 2016) or colour ornamentation (Griffith et al., 2003). Two of the focal populations studied here are located in habitats dominated by the evergreen holm oak (*Quercus ilex)* and the two others in habitats dominated by the deciduous downy oak (*Quercus humilis*). This environmental heterogeneity has been linked to strong phenotypic variation in blue tits, whereby birds from the two habitats differ in their morphology, behaviour, colour ornamentation, physiology and life histories (Blondel et al., 2006, Charmantier et al., 2016). Overall, the phenotypic divergence between the two types of habitats is consistent with a divergence in pace-of-life syndrome (Réale et al., 2010), with individuals in the deciduous habitat displaying a faster pace-of-life (e.g. larger clutches hence higher reproductive investment) and individuals in evergreen populations a slower pace-of-life (Charmantier et al., 2016, Dubuc-Messier et al., 2017). Such divergence in pace-of-life described on life-history and behavioural traits predicts lower adult survival associated with a faster pace-of-life syndrome in deciduous habitats.

Based on this context of a divergence in pace-of-life across habitats and on past investigations, we expected to find a difference in adult survival between habitat types (with lower survival probabilities in deciduous habitats), as well as between age classes (with lower survival in older individuals, Bouwhuis et al., 2012). We also expected year-to-year variation in adult survival to be correlated among populations as well as with global and local climatic indices (Grosbois et al., 2006, note their study is on three of the four populations studied here, and on a much shorter period of time (8 to 16 years) running until 2000). Finally, population density during the breeding season is also expected to have a negative impact on subsequent adult survival (e.g. Frederiksen & Bregnballe, 2000).

## Methods

### 1. Monitored populations

Data was collected in four Mediterranean wild populations of blue tits *Cyanistes caeruleus*, a European cavity-nesting non-migratory passerine bird, in forest plots equipped with nest boxes and dominated either by deciduous downy oaks (*Quercus pubescens*, site names starting with D-) or by evergreen holm oaks (*Quercus ilex*, site names starting with E-). All sites are situated in the Mediterranean region, three of them (E-Pirio: Lat 42.38; Long: 8.75; D-Muro: 42.55; 8.92; E-Muro: 42.59; 8.96) on the island of Corsica and one (D-Rouviere: Lat 43.66; Long 3.67) on the Mainland in southern France (Figure 1, Blondel et al., 2006, Charmantier et al., 2016 for details). All four populations were monitored as part of the same long-term research program, but monitoring did not begin the same year in each site (E-Pirio: 38 years of monitoring from 1979 to 2016, D-Rouviere: 26 years (1991-2016), D-Muro: 24 years (1993-2016), E-Muro: 19 years (1998-2016)).

**Figure 1:**
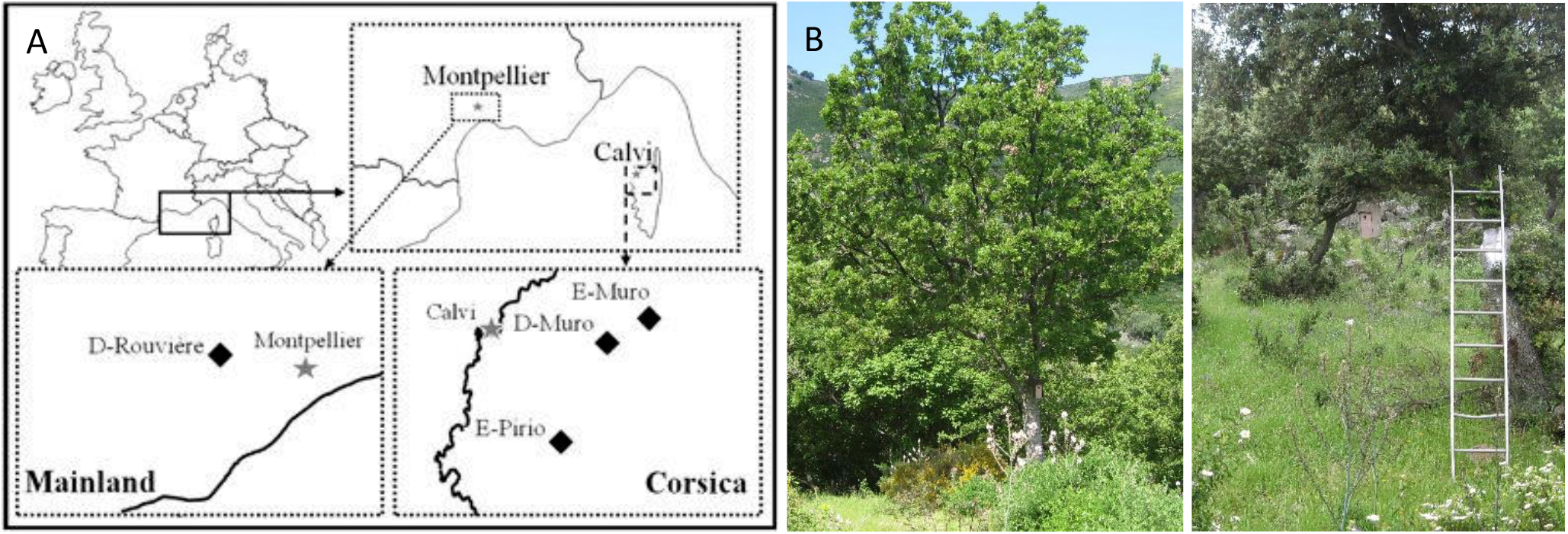
A/ Location of the four focal populations of blue tits. The three Corsican sites (E-Pirio, D-Muro and E-Muro) are located ~440km away from the mainland site (D-Rouviere). E-Muro is located ~6km away from D-Muro and ~30km from E-Pirio. Two sites are in a deciduous oak *Quercus pubescens* forest (D-Rouviere and D-Muro), and two in an evergreen oak *Quercus ilex* forest (E-Pirio and E-Muro). B/ A downy oak with a nest box in D-Muro, and C/ a holm oak with a nest box in E-Muro.

Populations were monitored using nest boxes, which blue tits readily use for breeding and generally prefer to natural cavities (Newton, 1994). The monitoring of almost all breeding individuals in the focal populations was ensured by using a high density of nest boxes compared to the abundance of natural cavities in the various sites (neighbouring boxes are 50m apart). In the four sites the total number of nest boxes varied across monitoring years: 103 to 234 boxes across 144 ha for D-Rouviere, 67-225 boxes across 108 ha for E-Pirio, 20-100 boxes across 45 ha for D-Muro and 20-76 nest boxes across 24 ha for E-Muro. In D-Rouviere, all nest boxes were wood-concrete Schwegler B1 boxes until 2012, with entrance hole diameters of either 28 mm or 32 mm. Since 2013, 15% of nest boxes in D-Rouviere were square layer larch boxes of three different sizes (see description of these wooden boxes in Lambrechts et al., 2017). Blue tit breeding densities were around 0.7 to 1.4 pairs/ha in D-Rouviere and E-Pirio, 0.8 to 1.9 pairs/ha in D-Muro, and 0.7 to 1.6 pairs/ha in E-Muro. In Corsica, nest predation was mainly attributed to green whip snakes, *Hierophis viridiflavus*, while in D-Rouviere, small mustelids such as the Least weasel *Mustela nivalis* or the Beech marten *Martes foina* are the main nestling predators. Over the years, several anti-predator devices were used such as placing nest boxes on posts rather than on trees, or placing the nest box in a wired cage. Note that while nest predation can be reported and reduced, predation events of adult individuals, e.g. by the Eurasian sparrowhawk *Accipiter nisus* remain unreported/unobserved, although they most probably represent an important mortality (Dhondt et al., 2008).

Every year, nest boxes were checked at least weekly during the blue tit breeding season (March to June). Adult breeders were captured in the boxes when the nestlings were 10-15 days old. The Centre de Recherches sur la Biologie des Populations d’Oiseaux (CRBPO) provided the permits under which capture and handling of birds were conducted (Permit n°1907 to Anne Charmantier as part of CRBPO capture program n°369), as well as unique numbered metal rings that were used to identify every (nestling and breeding) bird captured in the four populations. Plumage patterns were used to determine sex and age class at first capture (1 year breeders vs breeders of 2 or more years) for parents. Individuals whose minimum age at first capture had not been assessed were removed (3%, n=170 of parents). In total, 5499 reproducing individuals were considered in this study (E-Pirio: 1562, D-Rouviere: 1947, D-Muro: 1408, E-Muro: 582).

### 2. Large-scale climatic variation

Two large-scale climate indices were considered in this study as potentially influencing adult blue tit survival in our populations:

- The Mediterranean Oscillation Index (hereafter MOI, defined by Conte et al., 1989 and, Palutikof et al., 1996 as the normalized pressure difference between Algiers (36.4°N, 3.1°E) and Cairo (30.1°N, 31.4°E)). The MOI is a large scale climate index correlated to the North Atlantic Oscillation index (NAO) (Lionello et al., 2006). It is linked to rainfall and climate in the Mediterranean basin such that MOI is negatively correlated with precipitation and positively correlated with winter and spring temperature (Sangüesa-Barreda et al., 2019). Daily MOI data were obtained from https://crudata.uea.ac.uk/cru/data/moi/.
- The Sahel Rainfall index (monthly data obtained from http://research.jisao.washington.edu/data/sahel/), a tropical climate index.

The MOI was aggregated in two separate variables: winter MOI (from December 1^st^ year t to February 28^th^ in year t+1) and summer MOI (from June 15^th^ to September 15^th^ in year t). We initially intended to also include a yearly aggregation (from March 2^nd^ year t to March 1^st^ year t+1) yet it was strongly correlated to winter MOI (Pearson’s correlation coefficient = 0.857) hence dropped. The Sahel Rainfall index was aggregated in an yearly summer Sahel rainfall index (hereafter SRF) by adding the monthly values of June and July of each year, as has been done by Grosbois and colleagues on three of these blue tit populations (Grosbois et al., 2006).

### 3. Local meteorological data

Four local climatic variables were used in this study to test whether meteorological conditions drive differences in survival probability variability in each population:

- Spring and summer rainfall (hereafter SpringRF: aggregated rainfall between March 2^nd^ and September 15^th^ of year t)
- Autumn and winter rainfall (hereafter AutumnRF: aggregated rainfall between September 16^th^ of year t to March 1^st^ of year t+1)
- Extreme heat events during the summer (hereafter EHE, number of extremely hot days (average daily temperature in the 5% hottest of average daily temperatures in the studied summers on this site) between June 15^th^ and September 15^th^ of year t), which indicates if extreme climatic situations have been encountered during the considered summer
- Hottest summer temperatures (hereafter HST, average mean daily temperature during the 10 hottest days between June 15^th^ and September 15^th^ of year t), which represents the global harshness of the considered summer

Rainfall data were obtained by measurements from the meteorological station of Saint-Martin de Londres (Lat: 43.78 Long: 3.73, approximately 14 km away from the D-Rouviere site) for the mainland, and Calvi (Lat: 42.52; Long: 8.79, approximately 16 km away from E-Pirio, 11 km from D-Muro and 16 km from E-Muro) for the Corsican sites. Temperature data at the Corsican sites were obtained by regressing daily temperature measurements in the different sites with temperature data from the meteorological station of Calvi over 4 years, and inferring the temperature in the sites over the remaining years (see Table S1 in Appendix 1). At the D-Rouviere mainland site, temperature data was obtained from the meteorological station of Saint-Martin de Londres.

### 4. Population density index

Breeding density was obtained by measuring the blue tit nest box occupation rates in a restricted area within each study site. This area was defined based on two criteria: 1. It was located in the center of the full study area and 2. nest box locations and numbers were stable across study periods (E-Pirio: n=26 nest boxes in the first 6 years of monitoring (1979-1984), then 62 nest boxes, D-Rouviere: n=65 nest boxes, D-Muro: n=20 nest boxes the first year (1993), 38 boxes from 1997 to 1999 and 53 nest boxes in the remaining years, E-Muro: n=20 nest boxes during the first 3 years of monitoring (1998-2000), then 55 nest boxes).

### 5. Capture-Mark-Recapture modelling

Individual capture-recapture histories for the 5499 breeding individuals were analysed to provide robust estimates of survival and recapture probabilities (respectively ϕ and P, Lebreton et al., 1992), using a logit-link function. All analyses were conducted using the program E-SURGE (Choquet et al., 2009b). Goodness-of-fit of models to the data was ensured for each dataset using the program U-CARE (Choquet et al., 2009a), based on the Cormack Jolly Seber model for monostate models.

In several years, experiments were conducted in the population including brood size manipulations that can highly alter adult survival (Nur, 1984a, Dijkstra et al., 1990). Capture-recapture histories of the corresponding individuals were right-censored, after the first session they experienced such a fitness-changing experiment. However nestlings born during these experiments and later recaptured as adults were not removed from the analysis. This necessary censoring resulted in a significant decrease in observation numbers (−14.95% from 12 131 to 10 316 capture and recapture events in the dataset).

### 6. Model selection

The Akaike’s Information Criterion corrected for small sample size (AICc) was used for model selection (Burnham & Anderson, 1998). A low AICc was considered revealing a good compromise between the fit to the data (likelihood of the model) and the number of parameters used by the model. The threshold for a significant difference between two models was set at two AICc points. In case of a lower difference, the model with the lowest number of parameters was selected.

The first models (models integrating the four populations together, without temporal covariate) are numbered from 1 to 73 (Table 1). Covariate models (models integrating the four populations, with meteorological and population yearly covariates) are numbered 74 to 79 (Table 3). Finally, additional models were implemented separately for each population (models 80 to 111, Table 4).

**Table 1:** General model selection for models on pooled capture-recapture histories of the four focal blue tit populations, to test for the importance of age, sex, population (pop) and annual variation (year) on survival (ϕ) and recapture (P) probabilities. When a first order interaction between two variables is indicated, both non-interactive terms are present in the model even if not explicitly mentioned (e.g. Model 23: ϕ (pop.year) P (sex.pop) is equivalent to ϕ (pop + year + pop.year) P (sex + pop + sex.pop)). The best model is Model 1 with AICc = 10421.3358.

| Model<br>number | Model Description | | Number of | Deviance | $\Delta AICc$ |
| --- | --- | --- | --- | --- | --- |
| | $\phi$ | P | Parameters | | |
| 1 | $\phi$ (age + pop + year) | P (sex.pop) | 49 | 10320.75 | 0 |
| 2 | $\phi$ (sex + age + pop + year) | " | 50 | 10320.64 | 1.92 |
| 3 | $\phi$ (sex.age + pop + year) | " | 51 | 10319.66 | 2.96 |
| 4 | $\phi$ (age.pop + year) | " | 52 | 10319.73 | 5.06 |
| 5 | $\phi$ (sex.pop + age + year) | " | 53 | 10318.76 | 6.11 |
| 6 | $\phi$ (age.pop + sex + year) | " | 53 | 10319.58 | 6.93 |
| 7 | $\phi$ (sex.age + sex.pop + year) | " | 54 | 10318.06 | 7.44 |
| 8 | $\phi$ (sex.age + age.pop + year) | " | 54 | 10318.49 | 7.87 |
| 9 | $\phi$ (pop + year) | " | 48 | 10333.04 | 10.27 |
| 10 | $\phi$ (sex.pop + age.pop + year) | " | 56 | 10317.73 | 11.16 |
| 11 | $\phi$ (sex + pop + year) | " | 49 | 10333.04 | 12.29 |
| 13 | $\phi$ (sex.pop + year) | " | 52 | 10331.40 | 16.72 |
| 18 | $\phi$ (age + pop.year) | " | 112 | 10227.82 | 33.42 |
| 19 | $\phi$ (age + pop) | " | 13 | 10433.29 | 40.00 |
| 20 | $\phi$ (sex + age + pop) | " | 14 | 10433.17 | 41.89 |
| 21 | $\phi$ (age.pop) | " | 15 | 10432.08 | 42.81 |
| 22 | $\phi$ (sex.age + pop) | " | 15 | 10432.47 | 43.19 |
| 23 | $\phi$ (pop.year) | " | 111 | 10240.32 | 43.87 |
| 24 | $\phi$ (pop) | " | 12 | 10441.72 | 44.42 |
| 25 | $\phi$ (age.pop) | " | 17 | 10432.08 | 44.82 |
| 26 | $\phi$ (sex.pop + age) | " | 18 | 10431.53 | 46.27 |
| **27** | **$\phi$ (pop)** | **"** | **13** | **10441.72** | **46.43** |
| 28 | $\phi$ (age.pop + sex) | " | 18 | 10431.91 | 46.65 |
| 29 | $\phi$ (sex.age + sex.pop) | " | 19 | 10431.05 | 47.80 |
| 30 | $\phi$ (sex.age + age.pop) | " | 19 | 10431.11 | 47.86 |
| 32 | $\phi$ (sex + pop) | " | 14 | 10441.70 | 48.42 |
| 33 | $\phi$ (sex.pop) | " | 16 | 10440.24 | 50.97 |
| 34 | $\phi$ (sex.pop + age.pop) | " | 21 | 10430.32 | 51.09 |
| 35 | $\phi$ (sex.age + sex.pop + age.pop) | " | 22 | 10429.72 | 52.50 |
| 36 | $\phi$ (sex.pop) | " | 17 | 10440.24 | 52.98 |
| 37 | $\phi$ (age + year) | " | 47 | 10387.12 | 60.31 |
| 40 | $\phi$ (sex + age + year) | " | 48 | 10387.09 | 62.30 |
| 41 | $\phi$ (sex.age + year) | " | 49 | 10386.24 | 63.48 |
| **44** | **$\phi$ (year)** | **"** | **46** | **10396.34** | **67.51** |
| 45 | $\phi$ (sex + year) | " | 47 | 10396.34 | 69.53 |
| 48 | $\phi$ (age) | " | 11 | 10501.13 | 101.83 |
| 49 | $\phi$ (sex + age) | " | 12 | 10501.08 | 103.78 |
| 50 | $\phi$ (sex.age) | " | 13 | 10500.66 | 105.36 |
| **51** | **$\phi$ (.)** | **"** | **10** | **10507.34** | **106.03** |
| 54 | $\phi$ (sex) | " | 11 | 10507.34 | 108.03 |
| 12 | $\phi$ (sex.age + sex.pop + age.pop + year) | P (sex.pop) | 58 | 10316.91 | 12.36 |
| 14 | $\phi$ (sex.age + sex.pop + age.year) | " | 89 | 10268.19 | 26.71 |
| 15 | $\phi$ (sex.age + age.pop + age.year) | " | 89 | 10268.78 | 27.30 |
| 16 | $\phi$ (sex.pop + age.pop + age.year) | " | 91 | 10267.74 | 30.35 |
| 17 | $\phi$ (sex.age + sex.pop + age.pop + age.year) | P (sex.pop) | 92 | 10267.11 | 31.76 |
| 31 | $\phi$ (sex.age + sex.pop + age.pop + pop.year) | " | 121 | 10224.03 | 48.13 |
| 38 | $\phi$ (sex.age + sex.pop + age.year + pop.year) | " | 152 | 10172.66 | 60.76 |
| 39 | $\phi$ (sex.age + age.pop + age.year + pop.year) | " | 152 | 10172.66 | 60.76 |
| 42 | $\phi$ (sex.pop + age.pop + age.year + pop.year) | " | 154 | 10171.83 | 64.07 |
| 43 | $\phi$ (sex.age + sex.pop + age.pop + age.year + pop.year) | P (sex.pop) | 155 | 10171.21 | 65.53 |
| 46 | $\phi$ (sex.age + sex.pop + sex.year + age.pop + age.year) | " | 128 | 10236.91 | 75.42 |
| 47 | $\phi$ (sex.age + sex.pop + sex.year + age.pop + pop.year) | " | 157 | 10193.05 | 91.51 |
| 52 | $\phi$ (sex.age + sex.pop + sex.year + age.year + pop.year) | " | 188 | 10143.48 | 106.48 |
| 53 | $\phi$ (sex.age + sex.year + age.pop + age.year + pop.year) | " | 188 | 10144.03 | 107.03 |
| 55 | $\phi$ (sex.pop + sex.year + age.pop + age.year + pop.year) | " | 190 | 10142.76 | 109.94 |
| 56 | $\phi$ (sex.age + sex.pop + sex.year + age.pop + age.year + pop.year) | P (sex.pop) | 191 | 10142.01 | 111.23 |
| 57 | " | P (sex + pop) | 188 | 10156.10 | 119.10 |
| 58 | " | P (sex.pop + year) | 226 | 10084.81 | 127.56 |
| 59 | " | P (pop) | 187 | 10168.87 | 129.79 |
| 60 | " | P (sex) | 185 | 10177.07 | 133.80 |
| 61 | " | P (sex + year + pop) | 223 | 10098.43 | 134.86 |
| 62 | " | P (.) | 184 | 10189.13 | 143.78 |
| 63 | " | P (pop + year) | 222 | 10111.12 | 145.44 |
| 64 | " | P (sex + year) | 220 | 10119.32 | 149.43 |
| 65 | " | P (year) | 219 | 10127.43 | 155.44 |
| 66 | " | P (sex.pop + sex.year) | 260 | 10053.00 | 167.72 |
| 67 | " | P (sex.year + pop) | 257 | 10059.61 | 167.95 |
| 68 | " | P (sex.pop + pop.year) | 289 | 10000.48 | 177.04 |
| 69 | " | P (sex.year) | 254 | 10076.06 | 178.04 |
| 70 | " | P (pop.year + sex) | 286 | 10011.91 | 182.06 |
| 71 | " | P (pop.year) | 285 | 10021.25 | 189.26 |
| 72 | **$\phi$ (sex.age + sex.pop + sex.year + age.pop + age.year + pop.year)** | **P (sex.pop + sex.year + pop.year)** | **323** | **9962.89** | **212.50** |
| 73 | " | P (sex.year + pop.year) | 320 | 9971.25 | 214.39 |

A first model selection was conducted on the four populations together, assessing the effects of population, year, sex and age on recapture (*P*) and survival (ϕ) probabilities (Table 1). The starting model (Model 72) included potential effects of population, sex, age and annual variations (hereafter, *year*) on survival and potential effects of population, sex and year on recapture probabilities. Only simple (first order) interactions between variables influencing survival and recapture probabilities were considered, to enable robust biological interpretation of the results and avoid overcomplexity. All subsequent models were nested in this one. The model selection started with simplifying constraints on *P*. The best model structure for recapture (Model 56) was then retained and simpler survival models were considered, removing step by step first order interactions between variables until reaching a model with only 3 interactions (Model 12). All models nested in this one were then examined. At each step of model selection (selection on *P* structure, removal of interaction terms in ϕ, then selection of main effects in ϕ), new models were created and then sorted in descending AICc order. The model numbers in Table 1 reflect this step-by-step selection protocol.

To assess the potential effect of large-scale climatic fluctuations on survival in our populations, three large-scale climate indices (summer MOI, winter MOI, early summer Sahel Rainfall) were tested on the four populations together, assuming a linear or quadratic relationship with survival. As for models without covariates, model comparison was based on the AICc. Additionally, analyses of deviance (hereafter *ANODEV*, Skalski et al., 1993) were conducted to assess how much of the annual variation in survival could be explained by each candidate climate index. Each ANODEV used as general model the best model from the previous selection (Model 1) including additive effects of age, population and annual variations on survival, and as reduced model the same without the temporal effect (Model 19).

An analysis of deviance was also conducted to assess the effect of breeding density in year t on survival between t and t+1, in the different populations. In each population separately, the model including additive effects of age and year on survival was considered as the general model, while the reduced model included only an effect of age on survival. The same analysis was conducted in each population separately to assess the effect of local climatic variables (Table 4). A linear effect of time was also tested in this selection, to assess the existence of a potential temporal trend of survival probabilities in the different populations. All temporal covariates were centered and standardized.

## Results

### 1. Goodness of fit

No significant violation of standard model assumptions was found while performing the goodness-of-fit tests: for the 4-population monostate dataset χ^2^(df=602)=366.499, *p-value* = 1; for each population separately: χ^2^(df=184)=85.346, *p-value* = 1 for E-Pirio; χ^2^(df=140)=117.024, *p-value* = 0.922 for D-Rouviere; χ^2^(df=149)=101.227, *p-value* = 0.999 for D-Muro; χ^2^(df=129)=62.903, *p-value* = 1 for E-Muro.

### 2. Temporal variation in adult survival

Model selection indicated strong annual variation in survival probabilities across populations (Table 1, Figure 2). The effect of year was additive (see Table 1: Models 9, 23, 27, 44 and 51 for a comparison of additive and interactive effects of population and year on survival. ΔAICc(Model 9 – Model 23) = 33.59), which means that survival probabilities covaried across time in our 4 study populations (see Figure 2 for a visual testimony).

**Figure 2:**
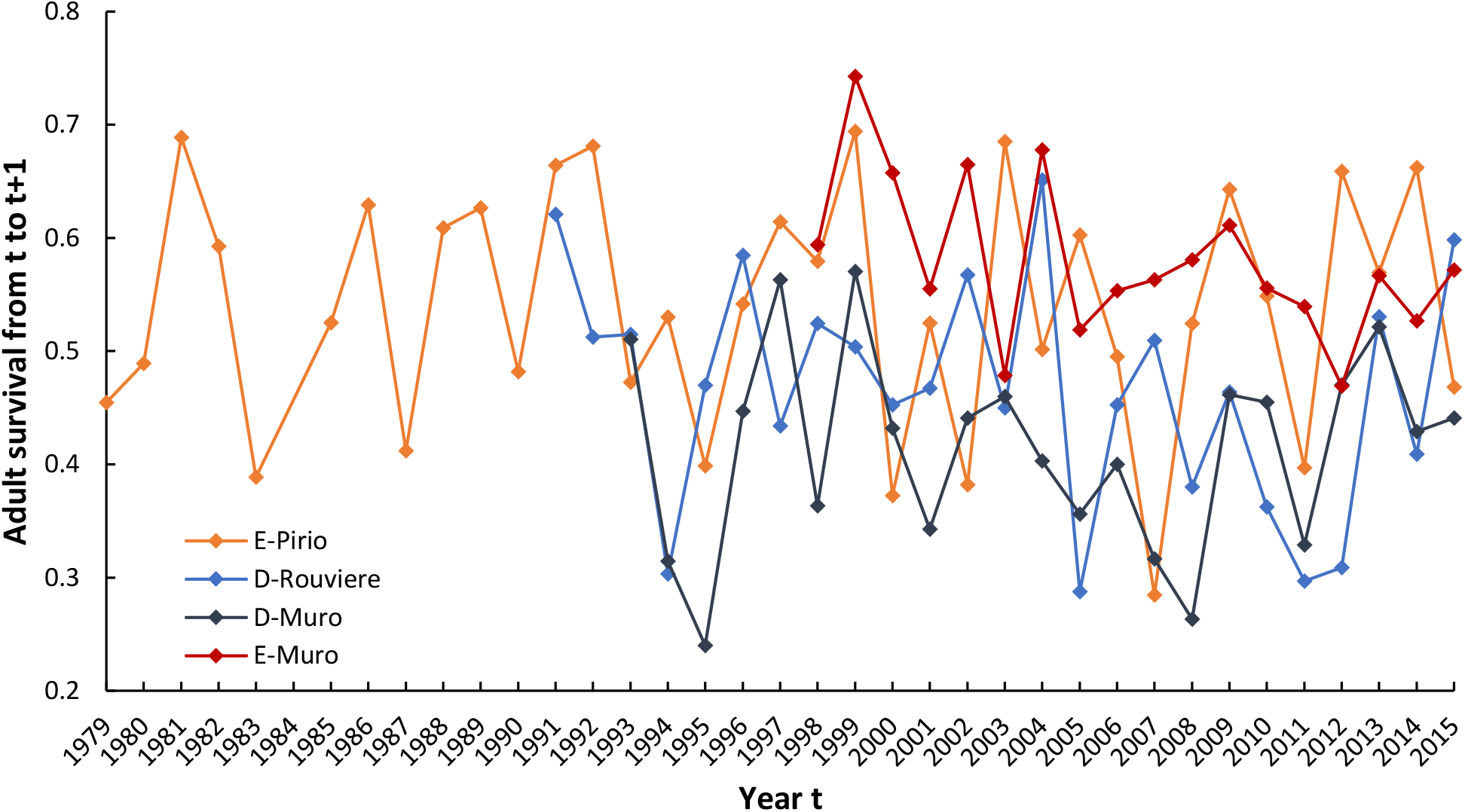
Annual survival for 2+ years adults in the four focal blue tit populations between breeding events in year t and t+1, from 1979 to 2016. 95% confidence intervals are not represented for the sake of readability. Estimates are from Model 18 with ϕ (age + pop.year) (Table 1).

### 3. Effects of population, age and sex on recapture and survival probabilities

Our model selection indicated that recapture probabilities differed across populations and sexes (Table 2) ranging from 0.673 (males in E-Pirio) to 0.866 (females in D-Rouviere). Additional model selection carried out separately for the four populations indicated that the effect of sex on P only occurred in E-Pirio (see Appendix 2 for additional model selection and Table 2 for estimates on P). In our best models, age had an additive impact on survival across populations, which means that while adult survival differed across populations, the effect of age on survival was similar across populations (Table 1). In particular, 1^st^ year adults had consistently markedly higher survival than adults aged 2 or more in all populations. In contrast, model comparison suggested that there was no effect of sex consistent in all populations (see model 1 vs. model 2 in Table 1) and the comparison of survival estimates for the two sexes with models not constrained to additivity indicated that the survival differences among sexes was weak and that the sex with the highest survival varied among populations (Table 2). Males had higher survival than females in deciduous habitats while the reverse was true in the evergreen populations (Table 2).

**Table 2:** Survival and recapture probability estimates and the influence of age and sex in the four focal populations of blue tits. 95% confidence intervals are enclosed in brackets.

|  | D-Rouviere | E-Pirio | D-Muro | E-Muro |
| --- | --- | --- | --- | --- |
|  | 26 years of monitoring<br>1947 individual histories | 38 years of monitoring<br>1562 individual histories | 24 years of monitoring<br>1408 individual histories | 19 years of monitoring<br>582 individual histories |
| **A) Annual Survival Estimates ($\phi$)** |  |  |  |  |
| **Adult survival ($\phi$)**<br>(estimates from Model 24) | 0.473<br>[0.453 , 0.493] | 0.548<br>[0.521 , 0.574] | 0.424<br>[0.397 , 0.451] | 0.575<br>[0.539 , 0.609] |
| **$\phi$ (1 year adults)**<br>(estimates from Model 21) | 0.491<br>[0.459 , 0.522] | 0.597<br>[0.547 , 0.645] | 0.449<br>[0.400 , 0.498] | 0.596<br>[0.522 , 0.666] |
| **$\phi$ (2+ years adults)**<br>(estimates from Model 21) | 0.458<br>[0.430 , 0.486] | 0.526<br>[0.494 , 0.558] | 0.412<br>[0.380 , 0.445] | 0.567<br>[0.525 , 0.608] |
| **$\phi$ (Males)**<br>(estimates from Model 33) | 0.479<br>[0.450 , 0.509] | 0.546<br>[0.508 , 0.583] | 0.429<br>[0.390 , 0.467] | 0.556<br>[0.505 , 0.606] |
| **$\phi$ (Females)**<br>(estimates from Model 33) | 0.467<br>[0.439 , 0.495] | 0.549<br>[0.513 , 0.585] | 0.419<br>[0.382 , 0.457] | 0.591<br>[0.543 , 0.638] |
| **B) Recapture Probability Estimates ($P$)** |  |  |  |  |
| **P (Males)**<br>(estimates from Model 1) | 0.821<br>[0.772 , 0.861] | 0.673<br>[0.611 , 0.729] | 0.749<br>[0.672 , 0.812] | 0.693<br>[0.606 , 0.768] |
| **P (Females)**<br>(estimates from Model 1) | 0.866<br>[0.819 , 0.902] | 0.857<br>[0.804 , 0.898] | 0.701<br>[0.625 , 0.767] | 0.740<br>[0.664 , 0.803] |

**Table 3:** Model selection procedure and significance of the effects of the considered climate indices on adult survival (ϕ). Starting yearly dependent and constrained model for the ANODEV are highlighted in bold. AICc of best model (Model 1) is 10421.34. SRF = Early Summer Sahel Rainfall; MOI= Mediterranean Oscillation Index. The values of R^2^ provided correspond to the proportion of temporal variance in survival explained by the model covariates, computed through analysis of deviance. The Beta column provides the estimate associated with each temporal covariate with its 95% confidence interval (for quadratic models, the two estimates correspond respectively to the linear and quadratic terms). Quadratic models are labelled with “q_”.

| Model | Model Description | Number of Parameters | Deviance | QAICc | ΔQAICc | Beta | p-ANODEV | R<sup>2</sup> |
| --- | --- | --- | --- | --- | --- | --- | --- | --- |
| 1 | **$\phi$ (age + pop + year) P (sex.pop)** | 49 | 10320.74 | 10421.33 | 0 |  |  |  |
| 74 | $\phi$ (age + pop + q_SRF) P (sex.pop) | 15 | 10423.86 | 10455.92 | 34.58 | 0.028 [-0.033;0.089]<br>-0.087 [-0.144;-0.030] | 0.21 | 0.084 |
| 75 | $\phi$ (age + pop + summerMOI) P (sex.pop) | 14 | 10427.56 | 10457.62 | 36.28 | -0.064 [-0.117;-0.012] | 0.17 | 0.051 |
| 76 | $\phi$ (age + pop + q_summerMOI) P (sex.pop) | 15 | 10427.26 | 10459.32 | 37.98 | -0.063 [-0.116;-0.010]<br>0.012 [-0.030;0.053] | 0.38 | 0.054 |
| 19 | **$\phi$ (age + pop) P (sex.pop)** | 13 | 10433.28 | 10461.33 | 39.99 |  |  |  |
| 77 | $\phi$ (age + pop + SRF) P (sex.pop) | 14 | 10432.81 | 10462.86 | 41.52 | -0.019 [-0.072;0.034] | 0.70 | 0.004 |
| 78 | $\phi$ (age + pop + winterMOI) P (sex.pop) | 14 | 10433.21 | 10463.26 | 41.92 | 0.008 [-0.047;0.062] | 0.89 | 0.001 |
| 79 | $\phi$ (age + pop + q_winterMOI) P (sex.pop) | 15 | 10432.94 | 10465.01 | 43.67 | -0.002 [-0.069;0.064]<br>-0.010 [-0.051;0.030] | 0.95 | 0.003 |

**Table 4:**
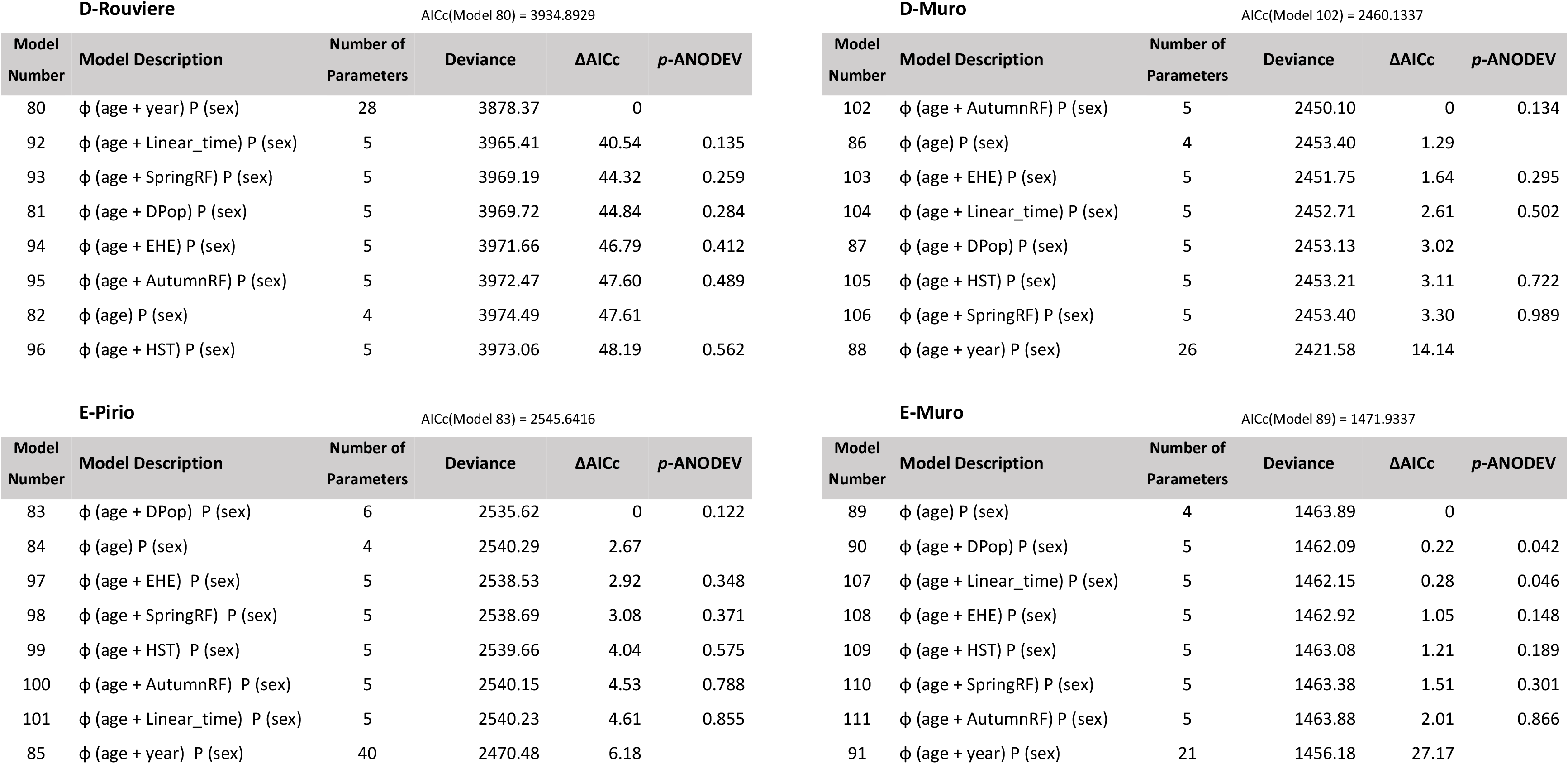
Effect of local climatic variables on adult survival in the four focal populations of blue tits. Dpop: breeding population density index, SpringRF : spring and summer rainfall, AutumnRF : autumn and winter rainfall, EHE : summer extreme heat events, HST: hottest summer temperatures, Linear_time: linear effect of time; see methods section for more details on these covariates.

| Model Number | Model Description | Number of Parameters | Deviance | ΔAICc | *p*-ANODEV |
| --- | --- | --- | --- | --- | --- |
| 80 | $\phi$ (age + year) P (sex) | 28 | 3878.37 | 0 | |
| 92 | $\phi$ (age + Linear_time) P (sex) | 5 | 3965.41 | 40.54 | 0.135 |
| 93 | $\phi$ (age + SpringRF) P (sex) | 5 | 3969.19 | 44.32 | 0.259 |
| 81 | $\phi$ (age + DPop) P (sex) | 5 | 3969.72 | 44.84 | 0.284 |
| 94 | $\phi$ (age + EHE) P (sex) | 5 | 3971.66 | 46.79 | 0.412 |
| 95 | $\phi$ (age + AutumnRF) P (sex) | 5 | 3972.47 | 47.60 | 0.489 |
| 82 | $\phi$ (age) P (sex) | 4 | 3974.49 | 47.61 | |
| 96 | $\phi$ (age + HST) P (sex) | 5 | 3973.06 | 48.19 | 0.562 |

**D-Muro** AICc(Model 102) = 2460.1337
| Model Number | Model Description | Number of Parameters | Deviance | ΔAICc | *p*-ANODEV |
| --- | --- | --- | --- | --- | --- |
| 102 | $\phi$ (age + AutumnRF) P (sex) | 5 | 2450.10 | 0 | 0.134 |
| 86 | $\phi$ (age) P (sex) | 4 | 2453.40 | 1.29 | |
| 103 | $\phi$ (age + EHE) P (sex) | 5 | 2451.75 | 1.64 | 0.295 |
| 104 | $\phi$ (age + Linear_time) P (sex) | 5 | 2452.71 | 2.61 | 0.502 |
| 87 | $\phi$ (age + DPop) P (sex) | 5 | 2453.13 | 3.02 | |
| 105 | $\phi$ (age + HST) P (sex) | 5 | 2453.21 | 3.11 | 0.722 |
| 106 | $\phi$ (age + SpringRF) P (sex) | 5 | 2453.40 | 3.30 | 0.989 |
| 88 | $\phi$ (age + year) P (sex) | 26 | 2421.58 | 14.14 | |

**E-Pirio** AICc(Model 83) = 2545.6416
| Model Number | Model Description | Number of Parameters | Deviance | ΔAICc | *p*-ANODEV |
| --- | --- | --- | --- | --- | --- |
| 83 | $\phi$ (age + DPop) P (sex) | 6 | 2535.62 | 0 | 0.122 |
| 84 | $\phi$ (age) P (sex) | 4 | 2540.29 | 2.67 | |
| 97 | $\phi$ (age + EHE) P (sex) | 5 | 2538.53 | 2.92 | 0.348 |
| 98 | $\phi$ (age + SpringRF) P (sex) | 5 | 2538.69 | 3.08 | 0.371 |
| 99 | $\phi$ (age + HST) P (sex) | 5 | 2539.66 | 4.04 | 0.575 |
| 100 | $\phi$ (age + AutumnRF) P (sex) | 5 | 2540.15 | 4.53 | 0.788 |
| 101 | $\phi$ (age + Linear_time) P (sex) | 5 | 2540.23 | 4.61 | 0.855 |
| 85 | $\phi$ (age + year) P (sex) | 40 | 2470.48 | 6.18 | |

**Table 4:** Effect of local climatic variables on adult survival in the four focal populations of blue tits.
| Model Number | Model Description | Number of Parameters | Deviance | ΔAICc | *p*-ANODEV |
| --- | --- | --- | --- | --- | --- |
| 89 | $\phi$ (age) P (sex) | 4 | 1463.89 | 0 | |
| 90 | $\phi$ (age + DPop) P (sex) | 5 | 1462.09 | 0.22 | 0.042 |
| 107 | $\phi$ (age + Linear_time) P (sex) | 5 | 1462.15 | 0.28 | 0.046 |
| 108 | $\phi$ (age + EHE) P (sex) | 5 | 1462.92 | 1.05 | 0.148 |
| 109 | $\phi$ (age + HST) P (sex) | 5 | 1463.08 | 1.21 | 0.189 |
| 110 | $\phi$ (age + SpringRF) P (sex) | 5 | 1463.38 | 1.51 | 0.301 |
| 111 | $\phi$ (age + AutumnRF) P (sex) | 5 | 1463.88 | 2.01 | 0.866 |
| 91 | $\phi$ (age + year) P (sex) | 21 | 1456.18 | 27.17 | |

### 4. Effect of large-scale climate indices on adult survival

Figure S1 (Appendix 3) presents temporal variation in the three considered large-scale climate indices (summerMOI, winterMOI and SRF) during the study period associated with models presented in Table 3. Among these covariates, only a small positive correlation was found between summerMOI and SRF (Pearson’s correlation coefficient_summerMOI-SRF_ = 0.362; *p-value* = 0.03). No correlation was found between winterMOI and the other climatic covariates. No significant linear nor quadratic effect of any of the tested large-scale climatic indices was found on survival probability (Table 3).

### 5. Effect of local meteorological variables on adult survival

Despite the apparent similar temporal variation across the four populations (Figure 2), the results of our model selection suggested important variation in adult survival across years only in the D-Rouviere population: ΔAICc = 47.61 between Model 80 and Model 82 (Figure 3B).

**Figure 3:**
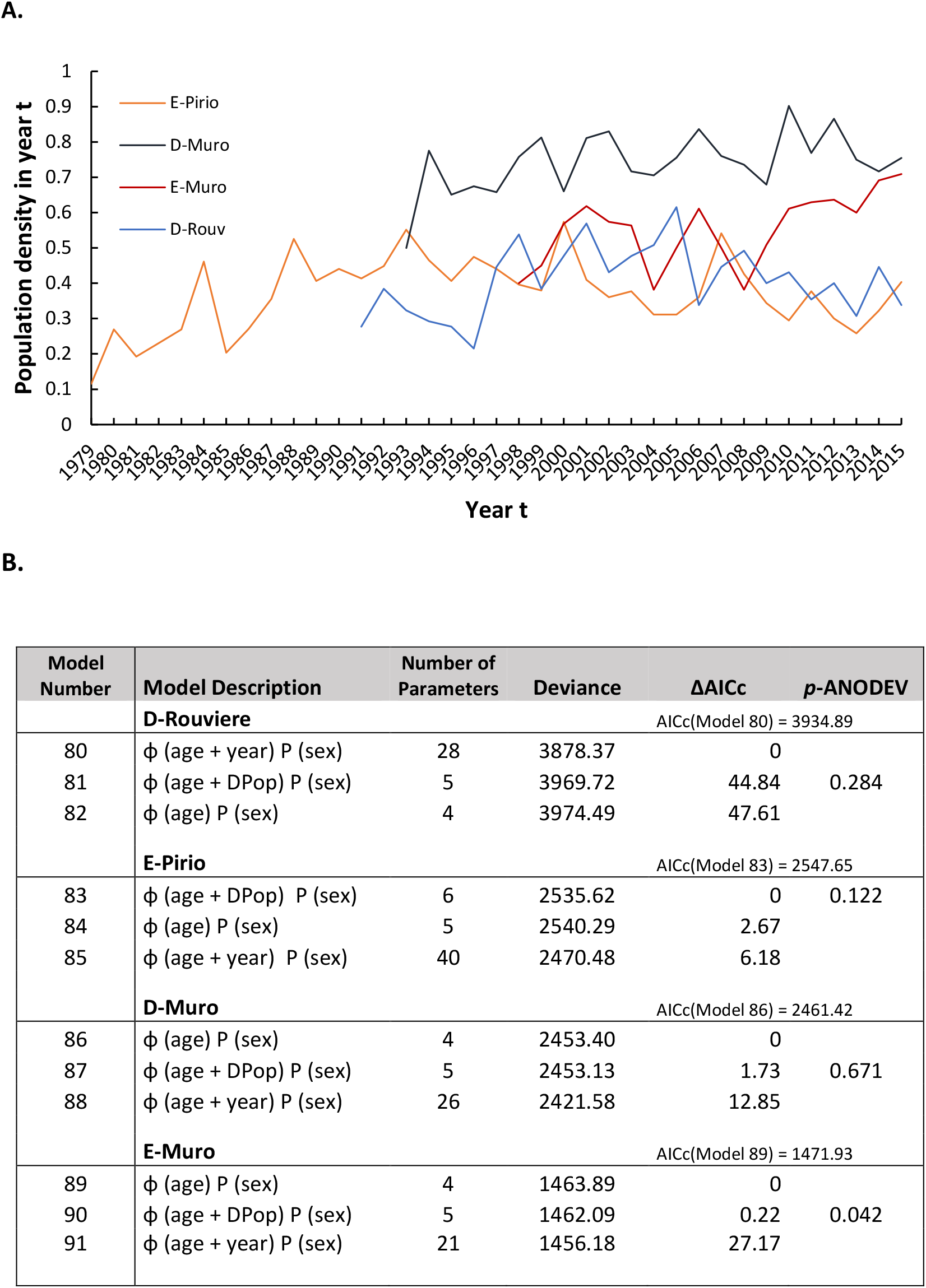
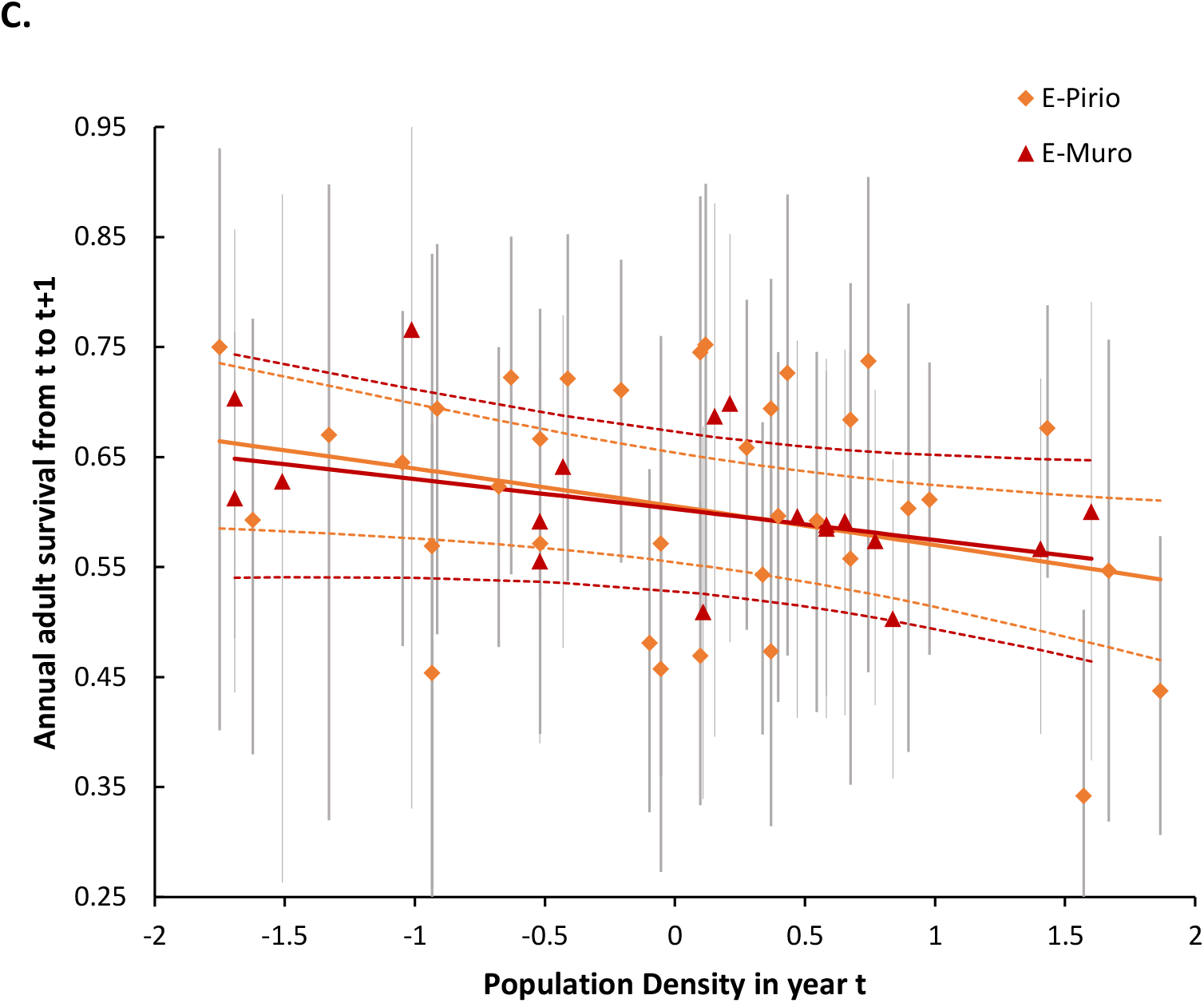
Effect of population density in year t on annual adult survival from t to t+1, in 4 populations of blue tits. A: Population density across the years of monitoring in the different populations. B: Model selection in each population separately to test for an effect of population density on adult survival. C: Annual adult survival as a function of population density in E-Pirio (orange) and E-Muro (red). Single points represent yearly estimates from Models 85 and 91 (see B) with 95% confidence intervals, lines represent the adult survival – population density relation as estimated by Models 83 and 90 (see B) with 95% confidence intervals in dotted lines.

To further explore these annual fluctuations in D-Rouviere and the difference with the other populations, we tested the effects of local weather variables (*i.e.* spring and summer rainfall, autumn and winter rainfall, extreme heat events, hottest summer temperatures, see Methods for details), which differed between the Corsican (E-Pirio, D-Muro, E-Muro) and the mainland (D-Rouviere) populations (Table 4). The existence of a linear temporal trend in survival was also tested in all four populations.

Within-population model selection provided support for a negative linear trend in survival in both D-Rouviere and E-Muro (Table 4). However, no effect of the tested meteorological variables was detected in any of the populations (Table 4).

### 6. Effect of breeding density on adult survival

We found no evidence that breeding densities were positively correlated across populations (Figure 3A): only one pairwise correlation was found whereby densities in E-Pirio and D-Muro were negatively correlated (correlation of −0.533, *p-value* = 0.007).

In one of the four populations (E-Pirio), the breeding density model had a lower AICc than both time-dependent and reduced models (ΔAICc was respectively 6.18 and 2.67, Figure 3B). For this population, the beta estimate associated with breeding density was significantly lower than zero (beta=−0.146, 95IC [−0.278; −0.013]) but the ANODEV test was not significant. In another population (E-Muro), the ANODEV test was slightly significant (*p*-ANODEV = 0.042) but the beta estimate was not significantly different from zero (beta=−0.116, 95IC [−0.286; 0.055]). In both cases, population density had a negative correlation with adult survival: in these two evergreen populations, springs with high nest box occupation were followed by a year of low adult survival (Figure 3C).

## Discussion

The present study provides estimates for adult survival and recapture probabilities in four Mediterranean blue tit populations breeding in nest boxes, and explores factors that explain variation in adult survival across years. The results show that these populations have different average survival probabilities, with higher survival in evergreen compared to deciduous forests, in accordance with the prediction of a slower pace-of-life in birds breeding in the less productive evergreen habitats. Despite these differences, the year-to-year variation in survival probabilities was similar among populations. This suggests that regional environmental conditions (e.g. climate) could drive survival fluctuations across populations. However, we did not find evidence that the atmospheric pressure conditions in the Mediterranean basin (approximated by the Mediterranean Oscillation Index MOI) nor the tropical climate index of rainfall in the Sahel were correlated to adult survival in these four blue tit populations. Estimates of breeding densities were not correlated across populations, but we found that in the two evergreen forests (E-Pirio and E-Muro) breeding density had a significant negative impact on adult survival during the following year. Finally, two of the focal populations (D-Rouviere and E-Muro) displayed a linear temporal decrease in adult survival over the study period.

### 1. Recapture probabilities across sex and population

Model selection for the global dataset indicated that sex had an effect on recapture probabilities and that this effect varied among populations. Further results (see Appendix 2 and Table 2) revealed that the effect of sex was significant only in the E-Pirio population, where females had higher recapture probabilities than males. In two other populations, D-Rouviere and E-Muro, recapture estimates were higher for females as well, but female and male confidence intervals largely overlapped (Table 2). Higher recapture probabilities for females may be explained either because they have higher provisioning rates than their male partners at the time when we capture parents in nest boxes (i.e. when nestlings are 10-15 days old, Banbura et al., 2001, Garcia-Navas et al., 2012, but see Limbourg et al., 2013, Iserbyt et al., 2019, and Garcia-Navas & Sanz, 2012 for evidence that sex-biased provisioning can shift across blue tit populations) or because males are more warry of human presence and traps used for nest box captures.

The fact that recapture probabilities are high in the D-Rouviere population (0.821 for males and 0.866 for females) is concordant with the scarcity of suitable cavities for breeding in the forest and the lower immigration rate in this population (Charmantier et al., 2014). Conversely, the three Corsican sites are surrounded by adequate forest patches that offer possibilities for blue tit natal or breeding dispersal outside the nest box area. Other factors potentially explaining different recapture probabilities across populations include differences in bird boldness or brood failure.

### 2. Survival probabilities across sex and population

The estimated survival probabilities in our different populations were comparable, yet on the upper range, of those previously published for blue tits further north of the species distribution (see Table 5 for published estimates of blue tit adult survival).

**Table 5:** A non-exhaustive list of estimates of blue tit annual adult survival previously published

| Blue tit adult survival estimates | Method | Location | Reference |
| --- | --- | --- | --- |
| *Males:* 0.17 - 0.31<br>*Females:* 0.19 - 0.41 | Not corrected by recapture probability: survival probably underestimated | Wytham Woods near Oxford, UK | (Nur, 1984b) |
| *Males:* 0.23 - 0.419<br>*Females:* 0.25 - 0.539 | Not corrected by recapture probability: survival probably underestimated | Four study plots near Antwerp, Belgium | (Dhondt, 1987) |
| *1st year adults:*<br>*Males:* $0.82 \pm 0.07$<br>*Females:* $0.65 \pm 0.09$ | Capture-mark-recapture (CMR) modelling | Ventoux forest, south of France | (Blondel et al., 1992) |
| *2+ year adults:*<br>*Males:* $0.68 \pm 0.05$<br>*Females:* $0.47 \pm 0.05$ | | | |
| *1st year adults (both sexes):*<br>$0.65 \pm 0.05$ | CMR modelling with data from 1979-1989 | E-Pirio, Corsica, France | (Blondel et al., 1992) |
| *2+ year adults (both sexes):*<br>$0.56 \pm 0.03$ | | | |
| *Males:* 0.30 - 0.63<br>*Females:* 0.26 - 0.59 | CMR modelling | Mixed boreal forest near Tammisaari, Finland | (Class et al., 2014) |
| *Males (unfaithful/faithful):* 0.31 / 0.51<br>*Females (unfaithful/faithful):* 0.26 / 0.48 | CMR modelling | Wytham Woods near Oxford, UK | (Culina et al., 2015) |
| *Males:* 0.429-0.556<br>*Females:* 0.419-0.591 | CMR modelling | D-Rouviere, mainland France & E-Pirio, D-Muro, E-Muro, Corsica | The present study |

Our results indicate that blue tit survival is higher in evergreen forest sites (E-Pirio and E-Muro) than in deciduous sites (D-Rouviere and D-Muro). One of reasons behind this difference between habitats might be that the permanent leaves of the evergreen forest better protect birds from aerial predators incurring extrinsic mortality. This result also fits with other facets of the phenotypic variation among these two habitats (morphology, behaviour, colour ornamentation, physiology and life history, Charmantier et al., 2016). Overall, the multivariate phenotypic differentiation documented between blue tits from deciduous and evergreen habitats is consistent with a divergence in pace-of-life syndrome (Réale et al., 2010) whereby individuals in the more productive deciduous habitat display a faster pace-of-life (in particular larger clutches and higher aggression) compared to individuals in evergreen populations (Dubuc-Messier et al., 2017, Charmantier et al., 2016). Our adult survival estimates are hence in agreement with this expectation (Table 2). A common garden experiment revealed that differences in behaviour (exploration speed and handling aggression) as well as physiology (heart rate) between D-Muro and E-Pirio were maintained when nestlings from the two areas were raised in aviaries and kept for 5 years (Dubuc‐Messier et al., 2018). This result suggests a genetic origin to the divergence in morphology and behaviour, yet survival in wild conditions could not be explored in this common garden experiment. The lower survival of blue tits in deciduous compared to evergreen habitats revealed in this study could hence result from a combination of differences in extrinsic (e.g. difference in predator or parasite rates) and intrinsic mortality. While the eco-evo determinants of differences in life histories, and more broadly pace-of-life across populations of the same species remain highly debated, even theoretically (Galipaud & Kokko, 2020, André & Rousset, 2020) we provide here an interesting case study whereby differences in survival are in line with the pace-of-life theory (Réale et al., 2010).

The CMR models also reveal an important effect of age, with adults of 2 years or more having a lower survival than 1st year breeders (Table 4). This result is in line with previous findings (see Table 5, Blondel et al., 1992) and might reveal actuarial senescence in blue tits. In a study of the closely related great tit over half a century in Wytham Woods (UK), survival probabilities were shown to rapidly decrease with age after two years (see Figure 1A in Bouwhuis et al., 2012). If age-specific survival follows a similar pattern in blue tits, aggregating survival estimates of all adults of 2 years or older would result in lower average survival for birds of 2 years or older than for one-year old birds.

### 3. Population density and survival

Breeding density was variable within populations with up to 30% increase or decrease between years in all populations (Figure 3A), in line with previous results (see Reed et al., 2013 for even stronger fluctuations in a Dutch Great tit population). A negative effect of population density on subsequent adult survival was expected as a consequence of the competition induced by high breeding density (Fay et al., 2015, Le Coeur et al., 2016), however it was only detected in the two evergreen populations (E-Pirio and E-Muro, Figure 3B&C) and density explained moderate amounts of temporal variance in survival in both cases (for E-Muro, R^2^=0.23, *p*-ANODEV = 0.042, for E-Pirio, R^2^=0.07, *p*-ANODEV = 0.12). This site-specific impact of breeding density on parental survival can be related to the higher environmental constraints in the less productive evergreen forests, where food resources are less abundant for blue tit nestlings (Blondel et al., 1991) and foraging distances higher for parents (Tremblay et al., 2005), suggesting higher breeding competition.

Comparing breeding density across populations of blue tits requires great caution. The density indices all describe year-to-year variation of nest box occupation in a portion of each focal population where nest box density is similar and stable over time. However, these occupations are most probably not driven by the same factors across populations, and in particular might be differentially related to intra- and inter-specific competition. As the number of nest boxes, their relative position, and the number of natural cavities vary between populations, a direct comparison is not easy to interpret. This limitation is a shortcoming of studying survival and recapture probabilities in nest box populations. While the type of data used here could not have been obtained without nest boxes which allow easy nest localisation, access to nestlings, and capture and recapture of parents, nest boxes also influence parent survival, e.g. by altering predation and parasitism rates (Burke et al., 2004), and by allowing for higher breeding densities.

### 4. Temporal variation in survival and the role of climate

Our results confirm previous findings based on shorter time scales (*i.e.* 10 annual estimations in E-Pirio in Blondel et al. (1992) and 6 to 14 annual estimates in Grosbois et al. (2006) for D-Muro, D-Rouviere and E-Pirio) whereby blue tit survival varies substantially across time and across populations. Exploring the drivers of temporal fluctuations however, gave quite contrasting results compared to previous findings. Indeed, contrarily to Grosbois et al. (2006), our longer-term analysis did not reveal any correlation between parental survival and large-scale climate indices, nor between survival and local meteorological variables. This major difference may have different origins: Grosbois et al. worked on one less population (E-Muro) and fewer years of monitoring (they focused on the 1985-2000 period) with a different data filtering. In particular, they did not right-censor adults that had undergone fitness-altering experiments such as brood-size manipulations, and it is possible that the birds undergoing experiments are sensitive to climatic conditions in a different/stronger way. Moreover, their model selection resulted in recapture probabilities differently constrained: contrarily to us, they allowed annual variations of recapture probabilities, which may have consequences on the estimated variation in survival. In order to understand the reasons for the discrepancy between our results and those of Grosbois et al. (2006), we conducted additional analyses on a restricted dataset (only including the populations and periods studied by Grosbois et al., see Table S3 in Appendix 4). The results indicated that as in the study by Grosbois et al., and contrary to our analysis of the full dataset, variation in survival was correlated with large-scale climatic variation. However, in our case, it was essentially the Mediterranean Oscillation Index (calculated over the summer period) that was correlated with adult tit survival, and not the Early Summer Sahel Rainfall. This suggests that the discrepancy between our results and those of Grosbois et al. (2006) concerning the link between survival and large-scale climatic variations is largely due to the dataset considered : our total sample covers 5499 life histories over periods of 19 to 38 years, while the reduced sample covers 1566 life histories over periods of 8 to 16 years. Overall, based on stronger datasets than previously used, we conclude that global and local climatic indices do not explain adult blue tit survival fluctuations over years.

We found that two populations (D-Rouviere and E-Muro) display a declining linear temporal trend in adult survival across 26 and 19 years respectively. Our analysis on small and large-scale climate variation suggests that this decline is not habitat-specific, and is not linked to direct (temperature, rainfall) meteorological causes, even though a significant warming has been reported in our study areas over the last three decades (Warming of 0.61-0.66 °C per decade in spring across the different populations, Bonamour et al., 2019). It might however be caused by indirect effects that are difficult to capture statistically, such as an influence of food availability (warming spring temperatures may cause a phenological mismatch between the blue tits and their prey, Visser & Gienapp, 2019, Visser et al., 1998), or of earlier blue tit phenology. In E-Pirio, springs with early blue tit breeding are followed by a year of low adult survival (Bastianelli et al., submitted work) hence it would be interesting to extend the meteorological analysis to conditions in spring at year t−1 influencing survival between year t and t+1. Overall, the present study does not provide evidence that the recent rapid climate change influences blue tit adult survival.

Although local and regional climate conditions were not identified as drivers in the temporal fluctuations in survival, an interesting result was that survival probabilities covaried across time in the 4 study populations (Model 1 in Table 1), consistently with Grosbois et al. (2006), even though temporal variability was of higher magnitude in the mainland population of D-Rouviere (Figure 2 and Table 3). The estimation of the inter-annual variance in survival depends on both sampling and process variances, which magnitudes are influenced by several parameters (e.g., sample size and proximity of mean survival to the 0 and 1 bounds). It is therefore complicated to formally compare the inter-annual variances obtained here with those from previous work. Nevertheless, year-to-year variation in survival of the order of 20-30% as observed here (Figure 2), is consistent with other studies in small passerines (see e.g. Siriwardena et al., 1999, Hõrak & Lebreton, 1998, Perdeck et al., 2000).

In conclusion, we have provided here estimations for survival probability of adult blue tits across four different populations, in two different habitats, thanks to four long-term monitoring projects. Parental survival from one spring to the next varied substantially across years, in a synchronous way across populations. Despite this synchrony, we found no evidence that climate is an important driver in this variation of adult survival, calling for further investigations in other mortality causes fluctuating in time, in particular resource abundance, predation and parasitism. We hope that this comparison across four populations that are relatively close (south of France) will inspire further comparisons at a larger scale (across the species distribution), including perhaps datasets where more biotic and abiotic environmental features have been monitored over time.

## Supporting information

Data_description

E-Surge_Models

Models_description

## Data accessibility

Two data files (CMR histories and covariates) and all models run in E-SURGE are shared as supplemental files in the bioRxiv submission of the manuscript.

## Acknowledgements

We are immensely grateful to all the field assistants, students, postdocs, technicians and researchers that have contributed to the long term monitoring of blue tits in Corsica and on the mainland. Special thanks to the long-term involvement of Jacques Blondel, Marcel Lambrechts, Philippe Perret, Arnaud Grégoire, Céline Teplitsky and Annick Lucas. We thank the PLT and SIE platform in CEFE for their help with fieldwork and data management. We thank Clotilde Biard, Aurélie Cohas, Pierre-Yves Henry, Nathalie Machon, Roger Pradel and Céline Teplitsky for useful discussions at several stages of the analyses. Long term funding support was received by the OSU-OREME. Version 4 of this preprint has been peer-reviewed and recommended by Peer Community In Ecology (https://doi.org/10.24072/pci.ecology.100085)

## Conflict of interest disclosure

The authors of this preprint declare that they have no financial conflict of interest with the content of this article.

## Supplementary material

## Appendix 1

**Table S1:**
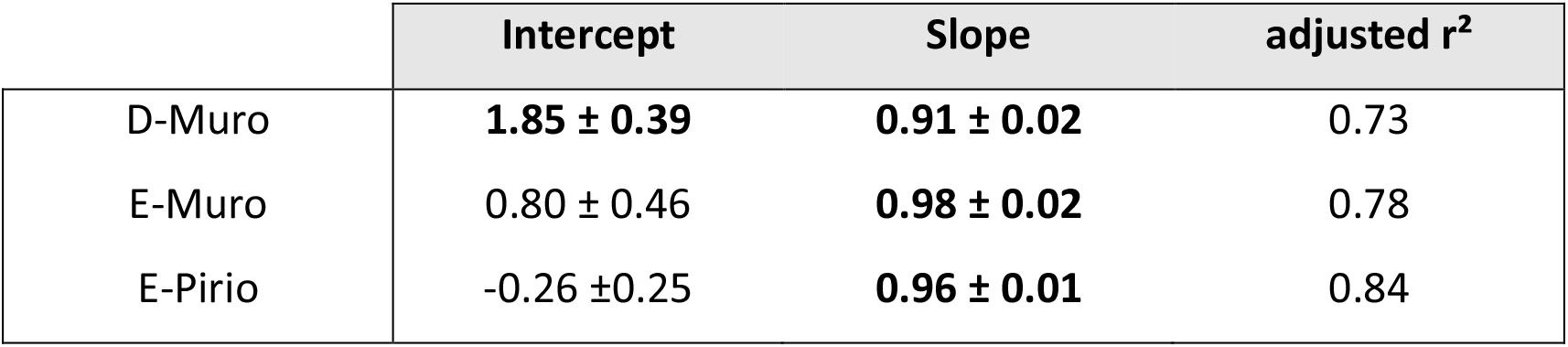
Correlation between local average daily temperature (minimum daily temperature + maximum daily temperature / 2) from thermocron i-buttons placed in trees near nest boxes, in the three Corsican sites (E-Pirio, E-Muro and D-Muro) and from measures of the national meteorological station of Calvi from 2013 to 2016. Significant estimates (p-value < 0.05) are highlighted in bold.

| | Intercept | Slope | adjusted $r^2$ |
| --- | --- | --- | --- |
| D-Muro | **$1.85 \pm 0.39$** | **$0.91 \pm 0.02$** | 0.73 |
| E-Muro | $0.80 \pm 0.46$ | **$0.98 \pm 0.02$** | 0.78 |
| E-Pirio | $-0.26 \pm 0.25$ | **$0.96 \pm 0.01$** | 0.84 |

## Appendix 2

In order to test *a posteriori* the effect of sex on the probability of recapture (P) in the study populations, we compared models with and without the effect of sex on P, considering for each population the survival structure of the best model (model 80 for D-Rouviere, model 84 for E-Pirio, model 86 for D-Muro, model 89 for E-Muro, see selection in Table 4 of the main manuscript).

The results presented below in Table S2 indicate that the effect of sex on the probability of recapture is significant only for the E-Pirio population.

**Table S2:** Comparison of models with and without a sex effect on recapture probabilities (P) in the four study populations of blue tits. In each population, model structure for survival (ϕ) is based on results from Table 4 of the main manuscript.

| Model description | Number of parameters | Deviance | AICc | $\Delta AICc$ |
| --- | --- | --- | --- | --- |
| **D-Rouviere** |  |  |  |  |
| $\phi(\text{age} + \text{year}) P(\text{sex})$ | 28 | 3878.37 | 3936.93 | 0 |
| $\phi(\text{age} + \text{year}) P(.)$ | 27 | 3880.46 | 3936.98 | 0.05 |
| **E-Pirio** |  |  |  |  |
| $\phi(\text{age}) P(\text{sex})$ | 4 | 2540.29 | 2550.32 | 0 |
| $\phi(\text{age}) P(.)$ | 3 | 2563.57 | 2571.59 | 21.27 |
| **D-Muro** |  |  |  |  |
| $\phi(\text{age}) P(.)$ | 3 | 2454.50 | 2462.52 | 0 |
| $\phi(\text{age}) P(\text{sex})$ | 4 | 2453.40 | 2463.43 | 0.91 |
| **E-Muro** |  |  |  |  |
| $\phi(\text{age}) P(.)$ | 3 | 1464.60 | 1472.63 | 0 |
| $\phi(\text{age}) P(\text{sex})$ | 4 | 1463.89 | 1473.95 | 1.31 |

## Appendix 3

**Figure S1:**
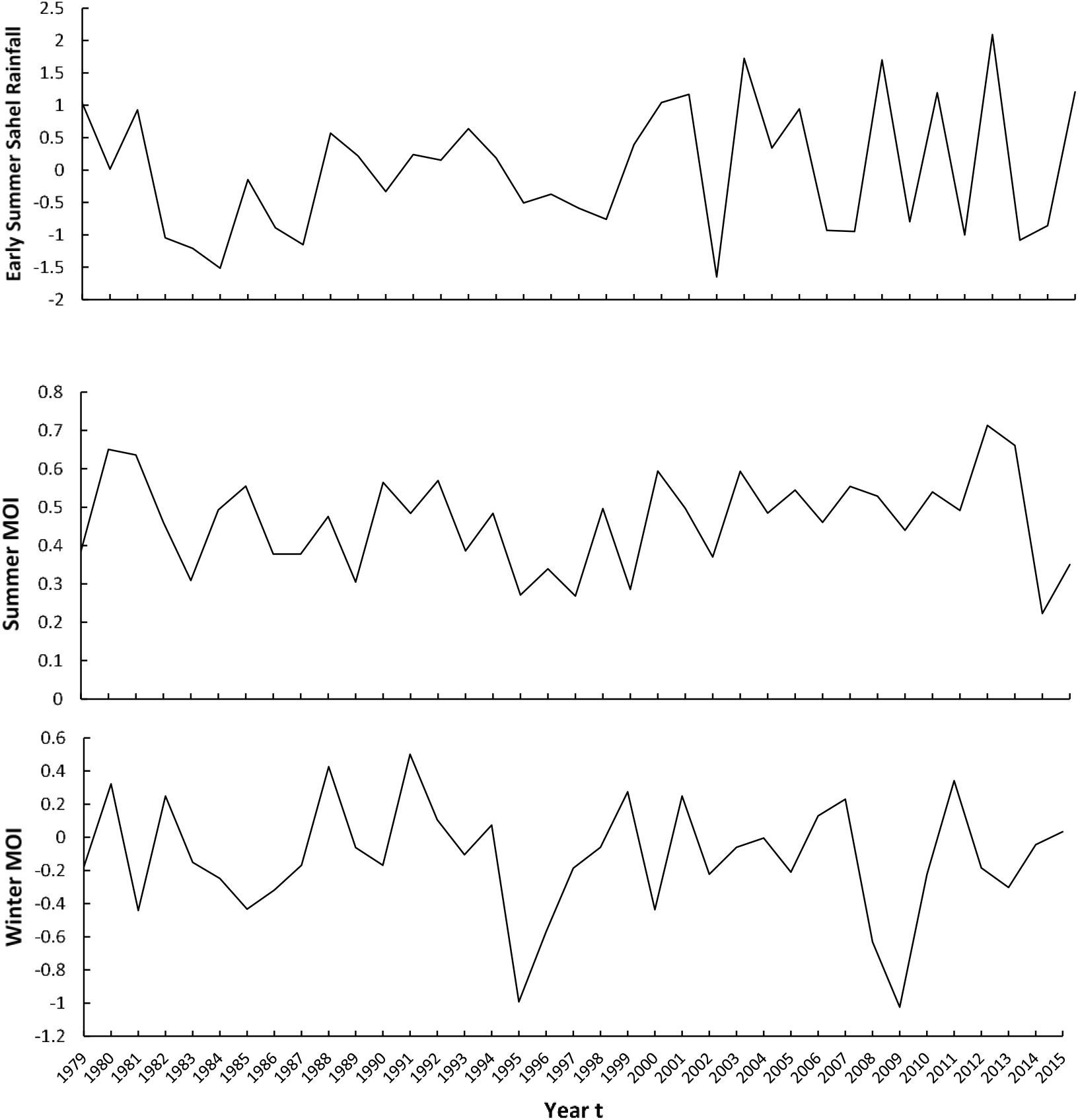
Annual variation in the three considered climate indices during the study period.

## Appendix 4

In order to understand the reasons for the discrepancy between our results and those of Grosbois et al. (Grosbois et al., 2006), who analyzed blue tit survival in three of our four study populations over a more limited period of time (8 to 16 years, datasets running until 2000), we restricted our dataset to years and populations studied by Grosbois and collaborators: E-Pirio : 1985-2000, D-Muro : 1993-2000 and D-Rouviere : 1991-2000. We then reanalysed the effect of large-scale climatic variation on mean survival observed in the three populations (based on the additive *Year*+*Pop* survival model).

The removal of the 1979-1984 and 2001-2016 periods and the E-Muro population restricted the dataset in terms of numbers of individuals (see Table S3 below, where our sample sizes are compared with those of Grosbois et al. (2006)).

**Table S3:** Sample sizes obtained with our restricted dataset and sample sizes used in Grobois et al. (2006).

| Study population | Nb. of individuals (restricted dataset, this study) | Nb. of individuals (Grosbois et al., 2006) |
| --- | --- | --- |
| D-Rouviere | 569 | 501 |
| E-Pirio | 689 | 692 |
| D-Muro | 308 | 253 |

To make our models comparable to those used by Grosbois et al. (2006), we used a general model ϕ (pop + year) P (sex + year.pop) and a reduced model ϕ (pop) P (sex + year.pop).

We tested the effects of three covariates (summerMOI, winterMOI and SRF) on survival variation through model comparison and ANODEV testing. Only linear effects of covariates were tested. The results are presented in Table S4 below (P-values do not include any correction for multi-testing).

The results indicate that as in Grosbois et al. (2006), and contrary to our analysis using a larger dataset, variation in survival is correlated with large-scale climatic variations. However, in our case, it was essentially the Mediterranean Oscillation Index (summerMOI, calculated over the summer period) that was correlated with adult tit survival, and not the Early Summer Sahel Rainfall (SRF). The discrepancy between these results is explained by the difference in datasets used.

**Table S4:** Model selection procedure and significance of the effects of three large-scale climatic indices on adult survival (ϕ) based on a restricted dataset (three populations over the period 1985-2000, see main text for details). Starting yearly dependant and constrained model for the ANODEV are highlighted in bold. AICc of the best model (first raw of the table) is 2449.71. SRF = Early Summer Sahel Rainfall; MOI= Mediterranean Oscillation Index. The values of R2 provided correspond to the proportion of temporal variance in survival explained by the model covariate, computed through an analysis of deviance. The Beta column provides the estimate associated with each temporal covariate with its 95% confidence interval.

| Model description | Nb.<br>parameters | Deviance | $\Delta AICc$ | Beta | p-ANODEV | R<sup>2</sup> |
| --- | --- | --- | --- | --- | --- | --- |
| $\phi(\text{pop} + \text{winterMOI}) P$<br>(sex+pop.year) | 37 | 2374.53 | 0 | 0.143<br>[0.021;0.265] | 0.023 | 0.284 |
| **$\phi(\text{pop}) P(\text{sex}+\text{pop}.\text{year})$** | 36 | 2379.73 | 3.14 |  |  |  |
| $\phi(\text{pop} + \text{SRF}) P(\text{sex}+\text{pop}.\text{year})$ | 37 | 2378.18 | 3.66 | 0.075<br>[-<br>0.043;0.193] | 0.27 | 0.084 |
| $\phi(\text{pop} + \text{summerMOI}) P$<br>(sex+pop.year) | 37 | 2379.71 | 5.19 | 0.010<br>[-<br>0.134;0.155] | 0.91 | 0.001 |
| **$\phi(\text{pop}+\text{year}) P(\text{sex}+\text{pop}.\text{year})$** | 50 | 2361.40 | 13.85 |  |  |  |
| $\phi(\text{pop}.\text{year}) P(\text{sex} + \text{pop}.\text{year})$ | 64 | 2351.36 | 33.18 | | | |

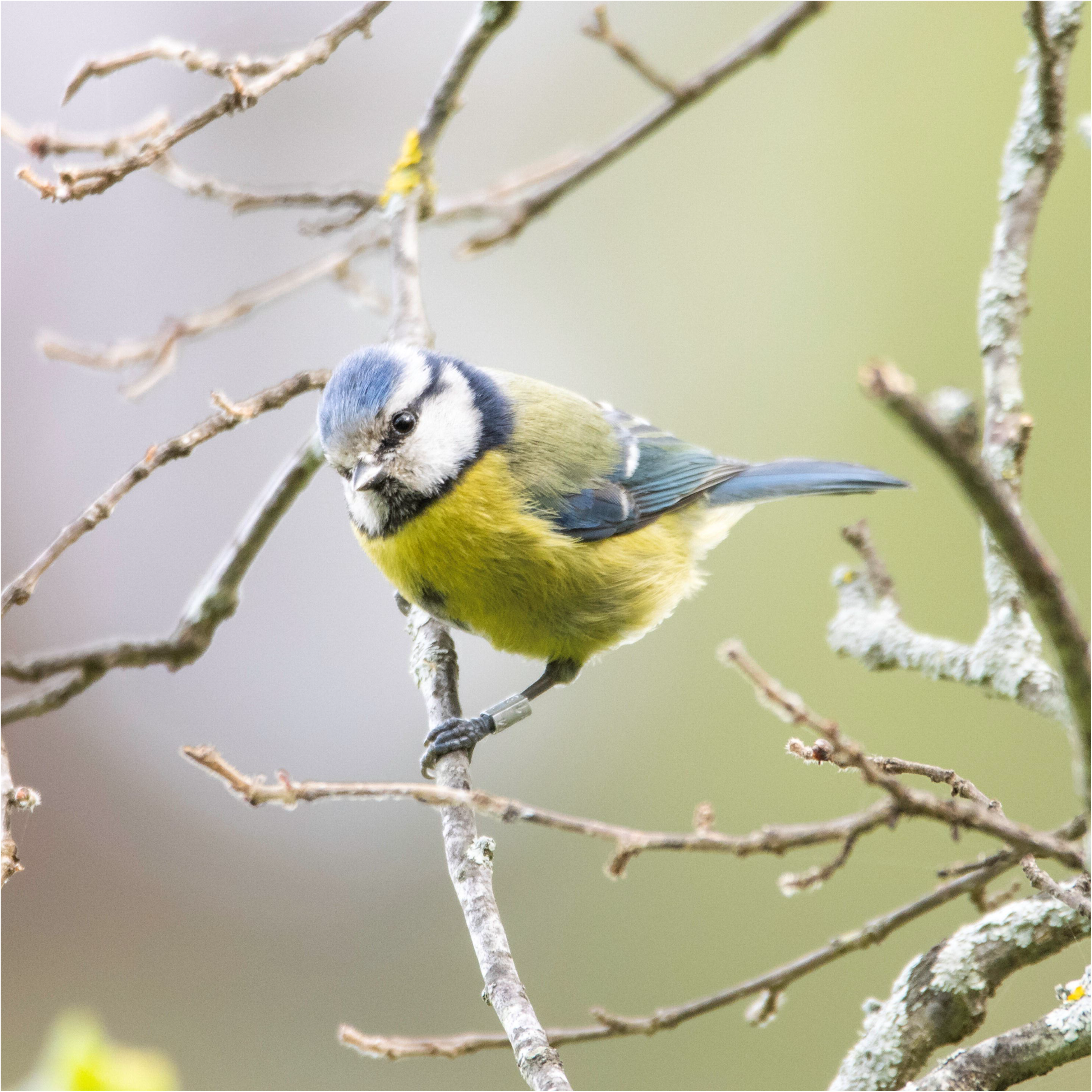

## Notes

### Competing Interest Statement

The authors have declared no competing interest.

### Summary of Updates

Version 4 of this preprint has been peer-reviewed and recommended by Peer Community In Ecology (https://doi.org/10.24072/pci.ecology.100085)

