## Supplementary material for "Identifying drivers of spatio-temporal variation in survival in four blue tit populations": Data_description

CMR DATA (CMR_histories.txt)

first column: CMR histories 1979-2016
second column: right censoring
third column: sex (1: male; 2: female)
fourth column: age class at first capture (1: 1 year old; 2: 2+ years old)
fifth column: site (population) (1=Rouviere, 2=Pirio, 3=D-Muro, 4=D-Muro)


TEMPORAL COVARIATES (Temporal_covariates.txt)

dpop PIR	:	population density in Pirio
dpop DMUR	:	population density in D-Muro
dpop EMUR	:	population density in E-Muro
dpop ROUV	:	population density in Rouviere
Ld PIR	:	Median laying date in Pirio
Ld DMUR	:	Median laying date in D-Muro
Ld EMUR	:	Median laying date in E-Muro
Ld ROUV	:	Median laying date in Rouviere
summerMOI	:	Mediterranean Oscillation Index (from June 15th to September 15th in year t)
winterMOI	:	Mediterranean Oscillation Index (from December 1st year t to February 28th in year t+1)
SuccesPir	:	Mean fledgling success in Pirio
SuccesDmur	:	Mean fledgling success in D-Muro
SuccesEmur	:	Mean fledgling success in E-Muro
SuccesRou	:	Mean fledgling success in Rouviere
TEPir	:	summer temperature: average daily average temperature from June 15 to September 15 year t  in Pirio
TEDmur	:	summer temperature: average daily average temperature from June 15 to September 15 year t  in D-Muro
TEEmur	:	summer temperature: average daily average temperature from June 15 to September 15 year t  in E-Muro
TERou	:	summer temperature: average daily average temperature from June 15 to September 15 year t  in Rouviere
THPir	:	 winter temperature: average of the daily average temperature from December 1st year t to Feb 28th year t+1  in Pirio
THDmur	:	 winter temperature: average of the daily average temperature from December 1st year t to Feb 28th year t+1  in D-Muro
THEmur	:	 winter temperature: average of the daily average temperature from December 1st year t to Feb 28th year t+1  in E-Muro
THRou	:	 winter temperature: average of the daily average temperature from December 1st year t to Feb 28th year t+1  in Rouviere
TCEPir	:	Summer warm temperature: average of the daily average temperature of the 10 warmest days between June 15 and September 15 of the year t  in Pirio
TCEDmur	:	Summer warm temperature: average of the daily average temperature of the 10 warmest days between June 15 and September 15 of the year t  in D-Muro
TCEEmur	:	Summer warm temperature: average of the daily average temperature of the 10 warmest days between June 15 and September 15 of the year t  in E-Muro
TCERou	:	Summer warm temperature: average of the daily average temperature of the 10 warmest days between June 15 and September 15 of the year t  in Rouviere
TFHPir	:	winter cold temperature: average of the daily average temperature of the 10 coldest days between December 1, year t and February 28, year t+1.  in Pirio
TFHDmur	:	winter cold temperature: average of the daily average temperature of the 10 coldest days between December 1, year t and February 28, year t+1.  in D-Muro
TFHEmur	:	winter cold temperature: average of the daily average temperature of the 10 coldest days between December 1, year t and February 28, year t+1.  in E-Muro
TFHRou	:	winter cold temperature: average of the daily average temperature of the 10 coldest days between December 1, year t and February 28, year t+1.  in Rouviere
PPPir	:	spring precipitation: average spring-summer year t: from March 2 year t to Sept. 15 year t  in Pirio
PPDmur	:	spring precipitation: average spring-summer year t: from March 2 year t to Sept. 15 year t  in D-Muro
PPEmur	:	spring precipitation: average spring-summer year t: from March 2 year t to Sept. 15 year t  in E-Muro
PPRou	:	spring precipitation: average spring-summer year t: from March 2 year t to Sept. 15 year t  in Rouviere
PAPir	:	fall precipitation: average autumn-winter years t-t+1: from 16 sept year t to 1 march year t+1  in Pirio
PADmur	:	fall precipitation: average autumn-winter years t-t+1: from 16 sept year t to 1 march year t+1  in D-Muro
PAEmur	:	fall precipitation: average autumn-winter years t-t+1: from 16 sept year t to 1 march year t+1  in E-Muro
PARou	:	fall precipitation: average autumn-winter years t-t+1: from 16 sept year t to 1 march year t+1  in Rouviere
EECPir	:	Extreme heat events: number of extremely hot days (average daily temperature in the hottest 5% over the monitoring period) from 15 June to 15 Sept year t  in Pirio
EECDmur	:	Extreme heat events: number of extremely hot days (average daily temperature in the hottest 5% over the monitoring period) from 15 June to 15 Sept year t  in D-Muro
EECEmur	:	Extreme heat events: number of extremely hot days (average daily temperature in the hottest 5% over the monitoring period) from 15 June to 15 Sept year t  in E-Muro
EECRou	:	Extreme heat events: number of extremely hot days (average daily temperature in the hottest 5% over the monitoring period) from 15 June to 15 Sept year t  in Rouviere
EEFPir	:	extreme cold events: number of extremely cold days from Dec. 1, year t to Feb. 28, year t+1.  in Pirio
EEFDmur	:	extreme cold events: number of extremely cold days from Dec. 1, year t to Feb. 28, year t+1.  in D-Muro
EEFEmur	:	extreme cold events: number of extremely cold days from Dec. 1, year t to Feb. 28, year t+1.  in E-Muro
EEFRou	:	extreme cold events: number of extremely cold days from Dec. 1, year t to Feb. 28, year t+1.  in Rouviere
EEPPir	:	extreme rainfall events: number of extremely wet days from spring t (March 2) to the end of winter t+1 (March 1)  in Pirio
EEPDmur	:	extreme rainfall events: number of extremely wet days from spring t (March 2) to the end of winter t+1 (March 1)  in D-Muro
EEPEmur	:	extreme rainfall events: number of extremely wet days from spring t (March 2) to the end of winter t+1 (March 1)  in E-Muro
EEPRou	:	extreme rainfall events: number of extremely wet days from spring t (March 2) to the end of winter t+1 (March 1)  in Rouviere
