## Supplementary material for "Identifying drivers of spatio-temporal variation in survival in four blue tit populations": Models_description

Abbreviations used in models'names

- phi : survival probability
- p : recapture probability
- i : initial state
- t : time (year effect)
- linear: linear effect of time
- a : age class
- captage : group of age class at first capture
- s : sex
- full : structure of the starting models for unspecified parameters
- CI : computation of confidence intervals


Abbreviations for covariates
popD: normalized population density in year t
EEC : number of extremely hot days (average daily temperature in the hottest 5% over the monitoring period) from 15 June to 15 Sept year t
TCE : average of the daily average temperature of the 10 warmest days between June 15 and September 15 of the year t
PP : spring precipitation: average spring-summer year t: from March 2 year t to Sept. 15 year t
PA : fall precipitation: average autumn-winter years t-t+1: from 16 sept year t to 1 march year t+1
sumMOI : Mediterranean Oscillation Index (from June 15th to September 15th in year t)
winMOI : Mediterranean Oscillation Index (from December 1st year t to February 28th in year t+1)
SFR : Sahel Rainfall index (addition of the monthly values of June and July of each year)
